# Karyotype asymmetry shapes diversity within the physaloids (Physalidinae, Physalideae, Solanaceae)

**DOI:** 10.1101/2020.04.21.053181

**Authors:** Julieta Rodríguez, Rocío Deanna, Franco Chiarini

**Affiliations:** Facultad de Ciencias Exactas, Físicas y Naturales (UNC), Vélez Sarsfield 299, Córdoba 5000, Argentina; Instituto Multidisciplinario de Biología Vegetal, IMBIV (CONICET-UNC), CC 495, Córdoba 5000, Argentina; Department of Ecology and Evolutionary Biology, University of Colorado, Boulder, CO, 80305, United States; Facultad de Ciencias Químicas (UNC), Medina Allende s.n., Córdoba 5000, Argentina

**Keywords:** Asymmetry, Chromosome evolution, Karyotype, Physalidinae, Solanaceae

## Abstract

Within the cosmopolitan family Solanaceae, Physalideae is the tribe with the highest generic diversity (30 genera and more than 200 species). This tribe embraces subtribe Physalidinae, in which positions of some genera are not entirely resolved. Chromosomes may help on this goal, by providing information on the processes underlying speciation. Thus, cytogenetic analyses were carried out in the subtribe in order to establish its chromosome number and morphology. Physalidinae is characterized by x = 12 and most species shows a highly asymmetric karyotype. These karyotype traits were mapped onto a molecular phylogeny to test the congruence between karyotype evolution and clade differentiation. A diploid ancestor was reconstructed for the subtribe, and five to six polyploidy independent events were estimated, plus one aneuploidy event (X = 11 in the monotypic genus *Quincula*). Comparative phylogenetic methods showed that asymmetry indices and chromosome arm ratio (r) have a high phylogenetic signal, whereas the number of telocentric and submetacentric chromosomes presented a conspicuous amount of changes. Karyotype asymmetry allow us to differentiate genera within the subtribe. Overall, our study suggests that Physalidineae diversification has been accompanied by karyotype changes, which can be applied to delimit genera within the group.

## 1. Introduction

Solanaceae, a cosmopolitan family of recognized economic and floristic importance, comprises 90–100 genera and 2400–3000 species (Olmstead & Bohs, 2006; Barboza et al.,2016). The tribe with the greatest number of genera is Physalideae, which includes more than 200 species (Olmstead et al., 2008). According to molecular phylogenetics studies (Olmstead et al., 2008; Särkinen, Bohs, Olmstead & Knapp, 2013), clades corresponding to the Iochrominae and Physalidinae subtribes of Physalideae have high supports (Olmstead et al., 2008; Särkinen et al., 2013), but the subtribe Withaninae and the subtribal position of some genera are not entirely resolved (Smith & Baum, 2006; Li, Gui, Xiong & Averett, 2013; Särkinen et al., 2013). Recently, a new phylogenetic hypothesis of Physalideae has been proposed, increasing sampling to 73 % of its species (Deanna, Larter, Barboza & Smith, 2019), which allows the analysis of evolutionary patterns and the proposal of taxonomic rearrangements.

Cytogenetics provides a valuable and irreplaceable source of information to address taxonomic, evolutionary and applied problems (Poggio, 1996; Guerra, 2012). Patterns and mechanisms of karyotypic evolution may explain diversification and speciation processes through the comparison of the chromosomes of different taxa (Weiss Schneeweiss & Schneeweiss, 2013). However, most of the cytogenetic information available from taxa belonging to Physalideae is restricted to reports of chromosome numbers and meiotic studies. This tribe has a basic number X = 12 (Badr, Khalifa, Aboel-Atta & Abou-El-Enain, 1997; Rego, da Silva, Torezan, Gaeta & Vanzela, 2009; Barboza, Chiarini & Stehmann, 2010; Chiarini, Moreno, Barboza & Bernardello, 2010; Deanna, Barboza & Scaldaferro, 2014), except for *Quincula* Raf. with X = 11 (Menzel, 1950). Many members of Physalideae have a meiotic chromosome number n = 12 (Moscone, 1992; Bohs, 2005; Sousa Peña, 2001), while *Withania* Pauquy, *Nothocestrum* A. Gray, *Tubocapsicum* Makino and some species of *Chamaesaracha* (A. Gray) Benth. and *Physalis* L. are polyploid, with meiotic numbers of either n = 24 or n = 36 (Menzel, 1950, 1951; Averett, 1973;Deanna et al., 2018).

Polyploidy is a common phenomenon in plants that occurs naturally and spontaneously, consisting of the increase in genome size caused by the presence of three or more chromosome sets (Otto & Whitton, 2000). This provides an important avenue for the evolution and generation of plant species (Winchester, 1981; Futuyma, 2005; Ranney, 2006; Thorpe, González Barrera & Rothstein, 2007; Hegarty & Hiscock, 2008; Maxime, 2008; Madlung, 2013). The ploidy level is an important factor to consider in crop improvement programs (Udall & Wendel, 2006). Indeed, there are studies on populations of cultivated cape gooseberry (*P. peruviana* L.) with 2n = 24, 36 and 48 (Rodríguez & Bueno, 2006). Chromosome counts in more Physalideae members are necessary to increase knowledge about the frequency and distribution of polyploidy events in the group.

The chromosome morphology (*i.e.*, karyotype) usually shows variability, mainly regarding to five different characteristics: (1) absolute size of the chromosomes, (2) centromere position, (3) relative chromosomes size, (4) basic number, and (5) number and position of the satellite (Stebbins, 1971). A symmetrical karyotype is one in which the chromosomes are approximately the same size and have a centromere which is in the middle position or slightly located towards one end. Increased asymmetry can occur through the change of the centromere position from the middle to the terminal position, or through the accumulation of differences in the relative size between the chromosome complement, thus yielding a more heterogeneous karyotype. These two trends are not necessarily correlated with each other, although they may be present in some groups (Stebbins, 1971). So far, karyoevolutive analyses in Physalideae have shown high variability in chromosome asymmetry between the subtribes, being the arm ratio (r) considered the chromosome trait with the greatest phylogenetic signal (*i.e.*, degree of similarity of characters associated with phylogenetic relationships between species), thus reflecting the effect of common ancestry (Deanna, Smith, Särkinen & Chiarini, 2018). Signal loss occurs when some species converge, in relation to a given trait, towards certain environmental conditions. The compartmentalization of processes that lead to the expression of traits reduces dependence between them, thus favouring their independent evolution (*i.e.*, closely related species do not tend to resemble each other; Deanna et al., 2018). However, in order to inquire about the evolutionary pattern and specifically about models of chromosome evolution within the subtribes, we need to increase cytogenetic analyses, especially for the Physalidinae subtribe for being the most widely distributed and the one with the greatest morphological diversity (Fig. 1) and specific richness (Zamora Tavares, Martínez, Magallón, Guzmán Dávalos & Vargas Ponce, 2016).

**Fig. 1.**
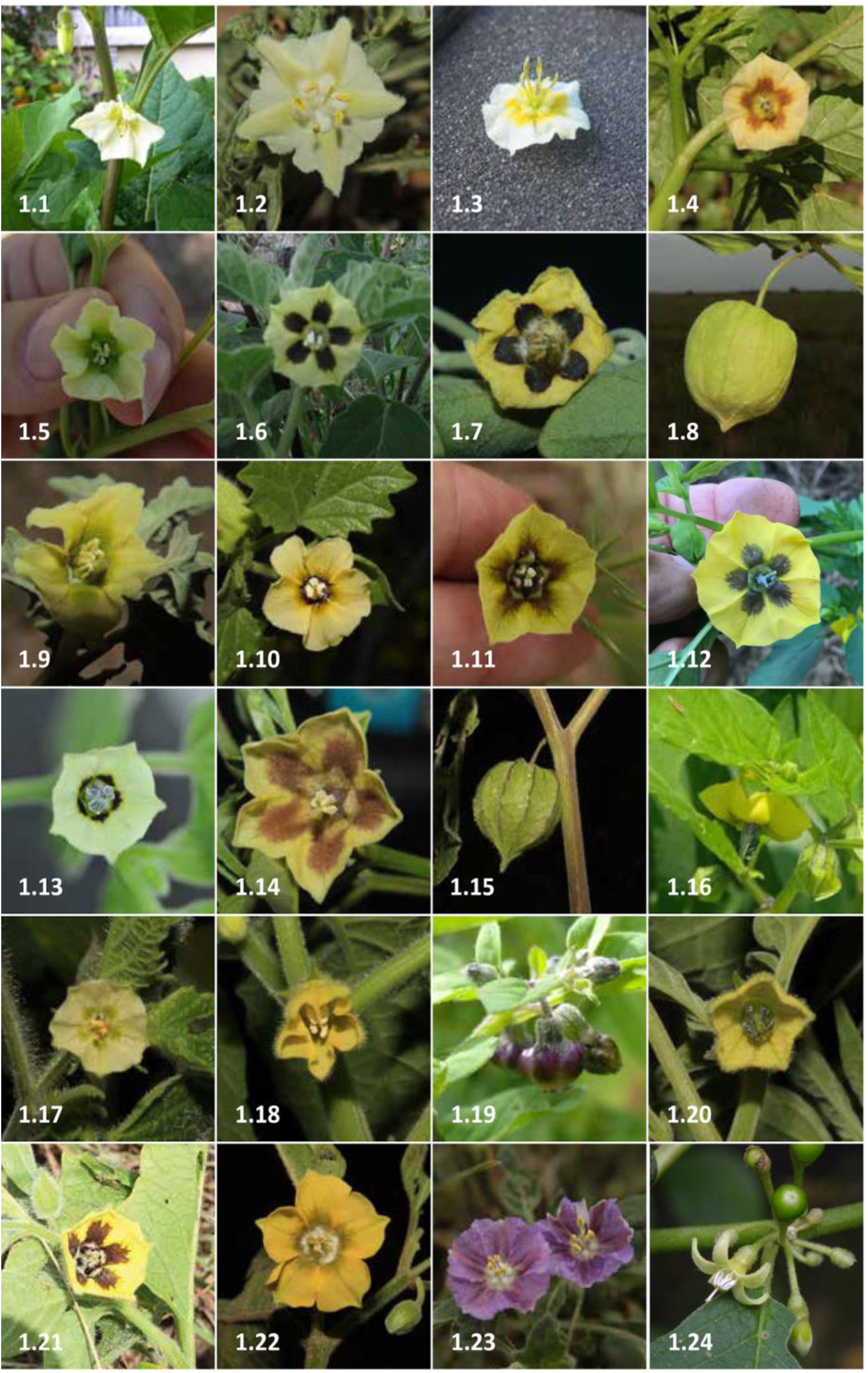
Species of Physalidinae currently cytogenetically analysed. (1.1*) Alkekengi officinarum* var. *officinarum*, flower (1.2) *C. coronopus*, fower (1.3*) P. acutifolia*, flower (1.4) *P. angulata*, flower (1.5) *P. crassifolia*, flower (1.6) *P. chenopodifolia*, flower (1.7*) P. cinerascens* var. *cinerascens*, flower (1.8) *P. cinerascens* var. *spathulifolia*, fruit (1.9) *P. fendleri*, flower (1.10) *P. hederifolia*, flower (1.11) *P. heterophylla*, flower (1.12) *P. ixocarpa*, flower (1.13) *P. lagascae*, flower (1.14*) P. longifolia*, flower (1.15) *P. peruviana*, fruit (1.16) *P. philadelphica*, flower (1.17) *P. pruinosa*, flower (1.18) *P. pubescens*, flower (1.19) *P. solanaceae*, mature fruits (1.20) *P. victoriana*, flower (1.21) *P. virginiana*, flower (1.22) *P. viscosa*, flower (1.23) *Q. lobata*, flower (1.24) *W. solanacea*, flower. Photographs by C. Pretz (3, 5, 16, 19); G. Chaniot (12); R. Deanna (2, 4, 6, 7, 8, 9, 10, 11, 14, 15, 17, 18, 20, 21, 22, 23, 24); S. Knapp (1); T. Sarkinen (13).

In the present study we aim to infer evolutionary patterns of chromosome characters onto a molecular phylogeny of the Physalidinae, accounting for an increase of karyological characterization of the genera within this subtribe, especially in the large genus *Physalis*. Regarding the current knowledge on ploidy level and how informative intrachromosomal and interchromosomal asymmetry have been to distinguish the subtribe Iochrominae (Deanna et al., 2018), we aim to characterize chromosomally the Physalidinae. Using comparative statistical methods, we address the following questions: (1) is there an increase of karyotype asymmetry in Physalidinae?, (2) are chromosome patterns informative enough to distinguish clades?, and 3) how many independent changes in the ploidy level have occurred in the group?

## 2. Materials and methods

### 2.1. Plant material

The provenance of the plant material used for cytogenetic studies is presented in Table 1. Seeds for this study were collected during field trips by F. Chiarini, G. E. Barboza and R. Deanna (Argentina, Colombia, Guatemala, Mexico, Peru and the United States). For each collection of ripe fruits, the location, geographical coordinates and date of collection were documented (Table. 1). Reference specimens were deposited in the herbarium of Museo Botánico de Córdoba (CORD), with duplicates in the herbaria of the province-country of material collection.

**Table 1.**
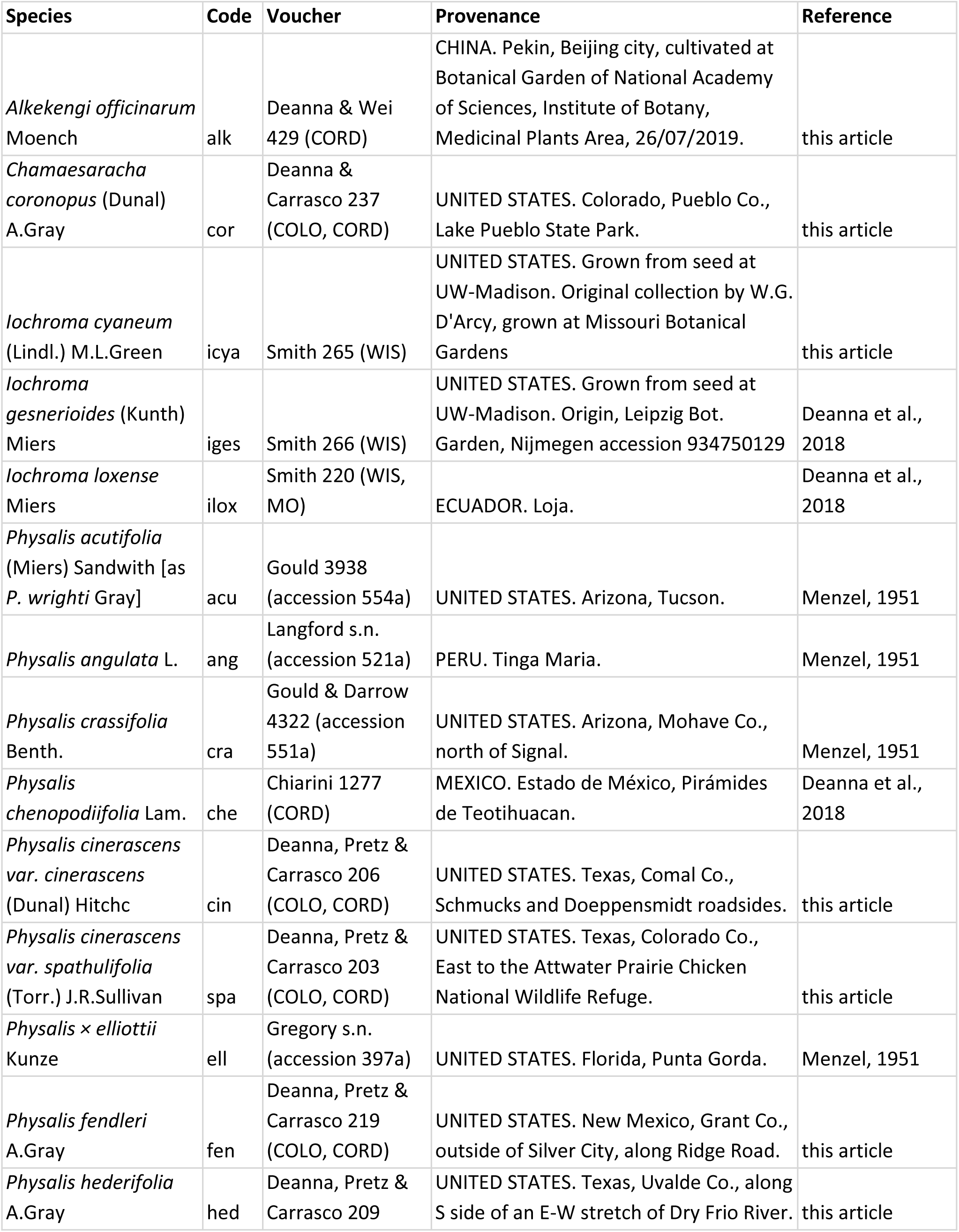
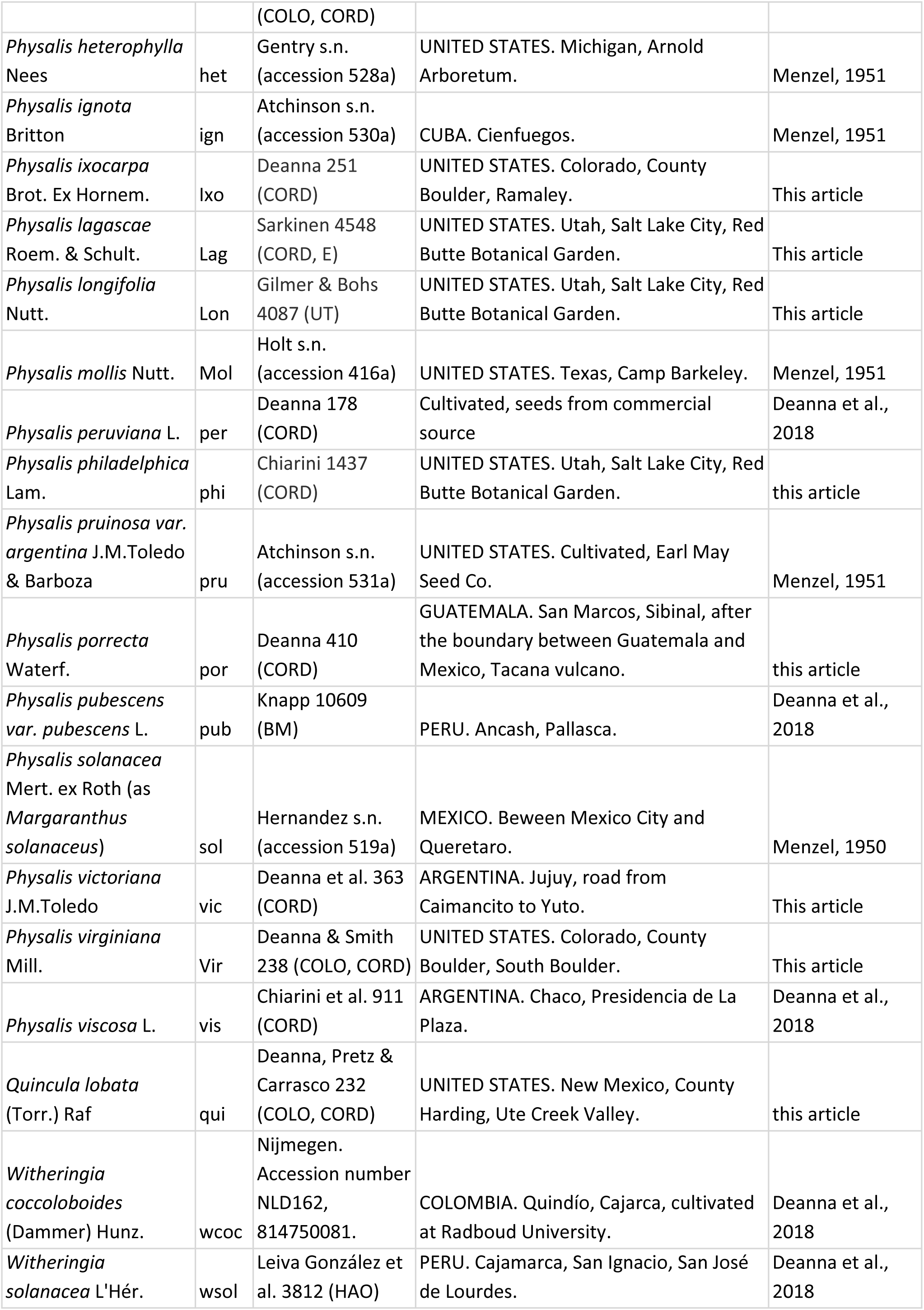
List of samples studied, code, provenance and reference (only if the species was previously studied) of all the analysed species of *Alkekengi*, *Chamaesaracha*, *Iochroma*, *Physalis*, *Quincula* and *Witheringia*.

| Species | Code | Voucher | Provenance | Reference |
| --- | --- | --- | --- | --- |
| *Alkekengi officinarum*<br>Moench | alk | Deanna & Wei<br>429 (CORD) | CHINA. Pekin, Beijing city, cultivated at Botanical Garden of National Academy of Sciences, Institute of Botany, Medicinal Plants Area, 26/07/2019. | this article |
| *Chamaesaracha coronopus* (Dunal)<br>A.Gray | cor | Deanna & Carrasco 237<br>(COLO, CORD) | UNITED STATES. Colorado, Pueblo Co., Lake Pueblo State Park. | this article |
| *lochroma cyaneum*<br>(Lindl.) M.L.Green | icya | Smith 265 (WIS) | UNITED STATES. Grown from seed at UW-Madison. Original collection by W.G. D'Arcy, grown at Missouri Botanical Gardens | this article |
| *lochroma gesnerioides* (Kunth)<br>Miers | iges | Smith 266 (WIS) | UNITED STATES. Grown from seed at UW-Madison. Origin, Leipzig Bot. Garden, Nijmegen accession 934750129 | Deanna et al., 2018 |
| *lochroma loxense*<br>Miers | ilox | Smith 220 (WIS, MO) | ECUADOR. Loja. | Deanna et al., 2018 |
| *Physalis acutifolia*<br>(Miers) Sandwith [as *P. wrightii* Gray] | acu | Gould 3938<br>(accession 554a) | UNITED STATES. Arizona, Tucson. | Menzel, 1951 |
| *Physalis angulata* L. | ang | Langford s.n.<br>(accession 521a) | PERU. Tinga Maria. | Menzel, 1951 |
| *Physalis crassifolia*<br>Benth. | cra | Gould & Darrow<br>4322 (accession 551a) | UNITED STATES. Arizona, Mohave Co., north of Signal. | Menzel, 1951 |
| *Physalis chenopodiifolia* Lam. | che | Chiarini 1277<br>(CORD) | MEXICO. Estado de México, Pirámides de Teotihuacan. | Deanna et al., 2018 |
| *Physalis cinerascens*<br>var. *cinerascens*<br>(Dunal) Hitchc | cin | Deanna, Pretz & Carrasco 206<br>(COLO, CORD) | UNITED STATES. Texas, Comal Co., Schmucks and Doeppensmidt roadsides. | this article |
| *Physalis cinerascens*<br>var. *spathulifolia*<br>(Torr.) J.R.Sullivan | spa | Deanna, Pretz & Carrasco 203<br>(COLO, CORD) | UNITED STATES. Texas, Colorado Co., East to the Attwater Prairie Chicken National Wildlife Refuge. | this article |
| *Physalis × elliotii*<br>Kunze | ell | Gregory s.n.<br>(accession 397a) | UNITED STATES. Florida, Punta Gorda. | Menzel, 1951 |
| *Physalis fendleri*<br>A.Gray | fen | Deanna, Pretz & Carrasco 219<br>(COLO, CORD) | UNITED STATES. New Mexico, Grant Co., outside of Silver City, along Ridge Road. | this article |
| *Physalis hederifolia*<br>A.Gray | hed | Deanna, Pretz & Carrasco 209 | UNITED STATES. Texas, Uvalde Co., along S side of an E-W stretch of Dry Frio River. | this article |
|  |  | (COLO, CORD) |  |  |
| *Physalis heterophylla* Nees | het | Gentry s.n. (accession 528a) | UNITED STATES. Michigan, Arnold Arboretum. | Menzel, 1951 |
| *Physalis ignota* Britton | ign | Atchinson s.n. (accession 530a) | CUBA. Cienfuegos. | Menzel, 1951 |
| *Physalis ixocarpa* Brot. Ex Hornem. | Ixo | Deanna 251 (CORD) | UNITED STATES. Colorado, County Boulder, Ramaley. | This article |
| *Physalis lagascae* Roem. & Schult. | Lag | Sarkinen 4548 (CORD, E) | UNITED STATES. Utah, Salt Lake City, Red Butte Botanical Garden. | This article |
| *Physalis longifolia* Nutt. | Lon | Gilmer & Bohs 4087 (UT) | UNITED STATES. Utah, Salt Lake City, Red Butte Botanical Garden. | This article |
| *Physalis mollis* Nutt. | Mol | Holt s.n. (accession 416a) | UNITED STATES. Texas, Camp Barkeley. | Menzel, 1951 |
| *Physalis peruviana* L. | per | Deanna 178 (CORD) | Cultivated, seeds from commercial source | Deanna et al., 2018 |
| *Physalis philadelphica* Lam. | phi | Chiarini 1437 (CORD) | UNITED STATES. Utah, Salt Lake City, Red Butte Botanical Garden. | this article |
| *Physalis pruinosa* var. *argentina* J.M.Toledo & Barboza | pru | Atchinson s.n. (accession 531a) | UNITED STATES. Cultivated, Earl May Seed Co. | Menzel, 1951 |
| *Physalis porrecta* Waterf. | por | Deanna 410 (CORD) | GUATEMALA. San Marcos, Sibinal, after the boundary between Guatemala and Mexico, Tacana vulcano. | this article |
| *Physalis pubescens* var. *pubescens* L. | pub | Knapp 10609 (BM) | PERU. Ancash, Pallasca. | Deanna et al., 2018 |
| *Physalis solanacea* Mert. ex Roth (as *Margaranthus solanaceus*) | sol | Hernandez s.n. (accession 519a) | MEXICO. Between Mexico City and Queretaro. | Menzel, 1950 |
| *Physalis victoriana* J.M.Toledo | vic | Deanna et al. 363 (CORD) | ARGENTINA. Jujuy, road from Caimancito to Yuto. | This article |
| *Physalis virginiana* Mill. | Vir | Deanna & Smith 238 (COLO, CORD) | UNITED STATES. Colorado, County Boulder, South Boulder. | This article |
| *Physalis viscosa* L. | vis | Chiarini et al. 911 (CORD) | ARGENTINA. Chaco, Presidencia de La Plaza. | Deanna et al., 2018 |
| *Quincula lobata* (Torr.) Raf | qui | Deanna, Pretz & Carrasco 232 (COLO, CORD) | UNITED STATES. New Mexico, County Harding, Ute Creek Valley. | this article |
| *Witheringia coccoloboides* (Dammer) Hunz. | wcoc | Nijmegen. Accession number NLD162, 814750081. | COLOMBIA. Quindío, Cajarca, cultivated at Radboud University. | Deanna et al., 2018 |
| *Witheringia solanacea* L'Hér. | wsol | Leiva González et al. 3812 (HAO) | PERU. Cajamarca, San Ignacio, San José de Lourdes. | Deanna et al., 2018 |

### 2.2. Classical staining and karyotype analyses

Seeds were germinated in Petri dishes under controlled photoperiod conditions (16: 8 h) and temperature (28 ± 0.5 ° C). To prevent germination failure, they were treated with gibberellic acid (GA_3_) at a concentration of 300 ppm (Ellis, Hong & Roberts, 1985). Slides with mitotic chromosomes were made from radical apices from germinating seeds treated with 8-hydroxyquinoline for 4 h, subsequently fixed in Farmer solution (3: 1 absolute ethyl alcohol and glacial acetic acid) at room temperature for 24 h and then stored at 4 ° C. The fixed roots were digested with a PECTINEX® enzyme solution (60 min at 37 ° C). Then, a conventional technique was applied for the observation of somatic chromosomes and their morphometric analyses, using Giemsa staining (Sheehan & Hrapchak, 1980).

At least 10 metaphases per sample were observed and micro-photographed with an Olympus BX61 optical microscope coupled to a monochromatic camera and Cytovision software (Leica Biosystems). Using the software ImageJ (Schneider, Rasband & Eliceiri, 2012), the following characteristics of each chromosome pair were recorded: s (short arm length), l (long arm length), c (total chromosome length) and satellite size. The relationship between the arms (r = l /s) was used to classify the chromosomes as metacentric (r = 1 to 1.7), submetacentric (r = 1.7 to 3), subtelocentric (r = 3 to 7), acrocentric (r = 7 to 8) and telocentric (r = 8 or more) according to the nomenclature of Levan, Fredga & Sandberg (1964). In addition, the haploid karyotype length (HKL), the mean chromosome length (c), the average of the relationship between arms or brachial index (r), and the ratio between the longest and the smallest chromosome in the complement (R) were calculated for each sample. The terminology of Battaglia (1955) was used to determine if the satellites correspond to microsatellites, macrosatellites or linear satellites. Idiograms based on the average values of each species or population were also made. Karyotype asymmetry was quantified according to Romero Zarco (1986) intrachromosomal (A_1_, which accounts for differences between arms of a single chromosome) and interchromosomal (A_2_, representing size differences among chromosomes of the same complement) asymmetry indices (Table 2).

**Table 2.** Chromosome data of karyotypically analysed species of Physalidinae: diploid number (2n); satellite pair order number (SAT); karyotype formula; total length of the karyotype haploid chromosome in μm ± standard deviation (HKL); total mean chromosome length in μm ± standard deviation (c); range of chromosome length (chr-l); mean arm ratio ± standard deviation (r); ratio between the longest and the shortest chromosome pair (R); intrachromosomal asymmetry index (A_1_); interchromosomal asymmetry index (A_2_).

| Species | 2n | SAT | HKF | HKL(sd) | c(sd) | chr-l | r(sd) | R | A1 | A2 |
| --- | --- | --- | --- | --- | --- | --- | --- | --- | --- | --- |
| *A. officinarum* var. *officinarum* | 24 | 1 | 9m + 3sm | 36.27 (4.11) | 3.02 (0.34) | 2.68 – 3.48 | 1.46 (0.06) | 1.30 | 0.28 | 0.12 |
| *C. coronopus* | 48 | 8 and 17 | 5m + 6sm + 1st | 53.23 (8.27) | 2.22 (0.35) | 1.62 – 2.94 | 2.06 (0.18) | 1.81 | 0.36 | 0.18 |
| *P. chenopodifolia* | 24 | 12 | 6m + 4sm + 2st | 38.05 (2.78) | 3.17 (0.23) | 2.80 – 3.99 | 1.87 (0.08) | 1.43 | 0.36 | 0.15 |
| *P. cinerascens* var. *cinerascens* | 24 | 9 | 8m + 3sm + 1st | 36.01 (4.66) | 3.00 (0.40) | 2.14 – 3.98 | 1.78 (0.28) | 1.86 | 0.33 | 0.20 |
| *P. cinerascens* var. *spathulifolia* | 24 | – | 7m + 4sm + 1st | 50.36 (3.01) | 4.20 (0.25) | 3.18 – 5.72 | 1.77 (0.25) | 1.80 | 0.34 | 0.21 |
| *P. fendleri* | 24 | 5 | 8m + 3sm + 1st | 35.46 (3.45) | 2.96 (0.29) | 2.28 – 3.74 | 1.67 (0.09) | 1.64 | 0.33 | 0.20 |
| *P. hederifolia* | 24 | 9 | 7m + 3sm + 2st | 37.61 (4.69) | 3.13 (0.39) | 2.07 – 4.07 | 1.92 (0.15) | 2.00 | 0.36 | 0.19 |
| *P. ixocarpa* | 24 | 4 | 6m + 5sm + 1st | 26.21 (3.46) | 2.18 (0.29) | 1.80 – 2.64 | 1.84 (0.16) | 1.47 | 0.39 | 0.16 |
| *P. lagascae* | 24 | – | 9sm + 3st | 29.99 (5.33) | 2.50 (0.44) | 2.06 – 3.00 | 2.77 (0.16) | 1.46 | 0.58 | 0.13 |
| *P. longifolia* | 24 | 8 | 3m + 7sm + 2st | 20.00 (5.02) | 1.67 (0.42) | 1.34 – 2.13 | 2.25 (0.52) | 1.60 | 0.49 | 0.16 |
| *P. peruviana* | 48 | 20 | 12m + 10sm + 1st + 1t | 48.26 (15.90) | 2.01 (0.66) | 1.08 – 2.58 | 2.23 (0.32) | 2.38 | 0.34 | 0.19 |
| *P. philadelphica* | 24 | 2 | 6m + 5sm + 1st | 31.05 (3.29) | 2.59 (0.27) | 2.16 – 3.17 | 1.81 (0.11) | 1.48 | 0.38 | 0.15 |
| *P. porrecta* | 24 | 5 | 5m + 5sm + 2st | 27.51 (5.41) | 2.29 (0.45) | 1.73 – 2.98 | 1.89 (0.28) | 1.72 | 0.40 | 0.20 |
| *P. pubescens* | 48 | 6 and 19 | 13m + 9sm + 1st + 1t | 60.27 (5.49) | 2.51 (0.23) | 1.65 – 3.26 | 2.36 (0.29) | 1.98 | 0.32 | 0.19 |
| *P. victoriana* | 24 | 10 | 5m + 6sm + 1st | 19.39 (1.94) | 1.62 (0.16) | 1.37 – 1.91 | 1.86 (0.22) | 1.40 | 0.40 | 0.12 |
| *P. virginiana* | 24 | 3 | 3m + 8sm + 1st | 23.87 (2.03) | 3.12 (0.17) | 1.90 – 2.56 | 2.02 (0.19) | 1.35 | 0.48 | 0.11 |
| *P. viscosa* | 24 | 11 | 6m + 5sm + 1st | 38.95 (3.36) | 3.25 (0.28) | 2.90 – 3.74 | 1.87 (0.11) | 1.29 | 0.40 | 0.14 |
| *Q. lobata* | 44 | – | 6m + 13sm + 3st | 44.91 (3.35) | 2.04 (0.15) | 1.84 – 3.16 | 2.08 (0.17) | 1.72 | 0.43 | 0.18 |

### 2.3. Chromosome characters database

A matrix was prepared with the chromosome continuous variables: c, r, A_1_ and A_2_. All the new measurements (except *P. porrecta*), and the species previously analysed by Menzel (1950, 1951; Table 1) were included. The amount of submetacentric and subtelocentric chromosomes was codified from the karyotype formula. Since these traits have a large number of discrete states, we codified them into three or less states. The amount of subtelocentric chromosomes was codified by establishing three states (‘0’ in case of absence of pairs, ‘1’ for species with a single pair and ‘2’ for species that contain two or more pairs of subtelocentric chromosomes). Similarly, submetacentric chromosomes were also codified in three states (‘0’ for species containing up to three pairs, ‘1’ for the ones that present four to six pairs and, finally. ‘2’ for species containing seven or more pairs).

### 2.4. Analysis of chromosome evolution in ChromEvol

Using the previously published Physalideae phylogeny (Deanna et al., 2019), chromosome number was reconstructed in ChromEvol (Mayrose, Barker & Otto, 2010; Glick & Mayrose, 2014), a software which was specifically developed to model the evolution of ploidy. RASP 3.2 program (Yu, Harris, Blair & He, 2015) was used to make inferences of the ancestral chromosome number in the subtribe, of the location of changes in chromosome number along the phylogeny, and of the total number of changes in the ploidy level of the group, such as it has been carried out in the subtribe Iochrominae (Deanna et al., 2018). All the models obtained were tested and compared in their probability values (AIC, Akaike, 1974). We set the base chromosome number as 12, the rate base number as 1, the maximal chromosome number as 120 (−10 according RASP settings), and the minimal chromosome number as 12 (1 according RASP settings). The base-number was kept fixed and 10,000 simulations were performed.

### 2.5. Ancestral reconstructions of chromosome characters

The evolution of discrete chromosome features was reconstructed on the maximum clade credibility (MCC) tree using the ace function of the package {ape} (Paradis, Claude & Strimmer, 2004) and the function make.simmap of the package {phytools} (Revell, 2012), in R version 3.4.2 (R Core Team, 2017). The ancestral states were inferred under a model where all the transition rates are different (“ARD”). Bayesian stochastic mapping (Nielsen, 2001; Huelsenbeck, Nielsen & Bollback, 2003), was performed with 1,000 simulations on the MCC tree obtained from Deanna et al. (2019). The reconstruction was applied to all species considering the unknown data as ambiguous and inferring the states of these species during reconstruction. We estimated the median number of changes per transition, generally preferred over means for non-normal distributions, and the 95% credibility interval using the hdr function of the package {diversitree} in R (FitzJohn, 2012).

The remaining cytogenetic characters (c, r, A_1_ and A_2_), coded as continuous, were mapped and plotted onto the MCC tree, after removing those species without data using the drop.tip function of the package {ape} in R (Paradis et al., 2004). The ancestral states were estimated assuming that the species evolve under a Brownian model and the mapping was performed using the ContMap function of the package {phytools} in R version 3.4.2 (Revell, 2012).

### 2.6. Phylogenetic signal of continuous traits

Reconstructions of ancestral states detect whether evolutionary shared stories, according to phylogeny, are the cause of the similarity patterns observed in the data. Therefore, the phylogenetic signal was evaluated based on the set of the continuous characters mentioned before. The statistics λ (Pagel, 1999) and K of Blomberg, Garland & Ives (2003) were inferred using the phylosig function of the package {phytools} in R (Revell, 2012). Values of λ close to zero correspond to traits that are less similar among species than expected for their phylogenetic relationships, so it follows that there is no phylogenetic signal. Otherwise, values of λ close to one correspond to traits that are more similar among closely related species, so it is estimated that there is a phylogenetic signal within the group. Likewise, the highest values of K indicate a high phylogenetic signal, with a value of one corresponding to the expected covariance under a Brownian evolution model. It was tested if K is significantly different from one when compared to inferred K values after 10,000 Brownian trait evolution simulations, implementing the fastBM function of the package {phytools} in R (Revell, 2012). We also verify if K is significantly different from zero (without phylogenetic signal) by comparing the estimated values of K with 10,000 null models where the species are randomly mixed using the phylosignal function of the {spicy} package in R (Kembel et al., 2010).

## 3. Results

Karyotype information was recorded for 42 species, including 18 new reports. Chromosome number, ploidy level and karyotype formula were analysed using classical cytogenetic techniques (Table 2, Fig. 1, Suppl. mat. S1).

### 3.1. Chromosome numbers and morphology

Physalidinae has a basic number X = 12, except *Quincula* with X = 11. Most species resulted diploid, except the species of the genus *Chamaesaracha*, *Leucophysalis grandiflora* and four species of *Physalis*, which are polyploid. Most species presented a single pair of chromosomes with satellites, except for the tetraploid species that presented two pairs and for *Quincula* which presented none. The satellites were always located in the short arm of one of the metacentric, submetacentric or subtelocentric pairs (Fig. 2; Suppl. mat. S1). Among the studied species that presented satellite, only five presented microsatellites and 10 presented macrosatellite. No species were observed with linear satellites (Suppl. mat. S1).

**Fig. 2.**
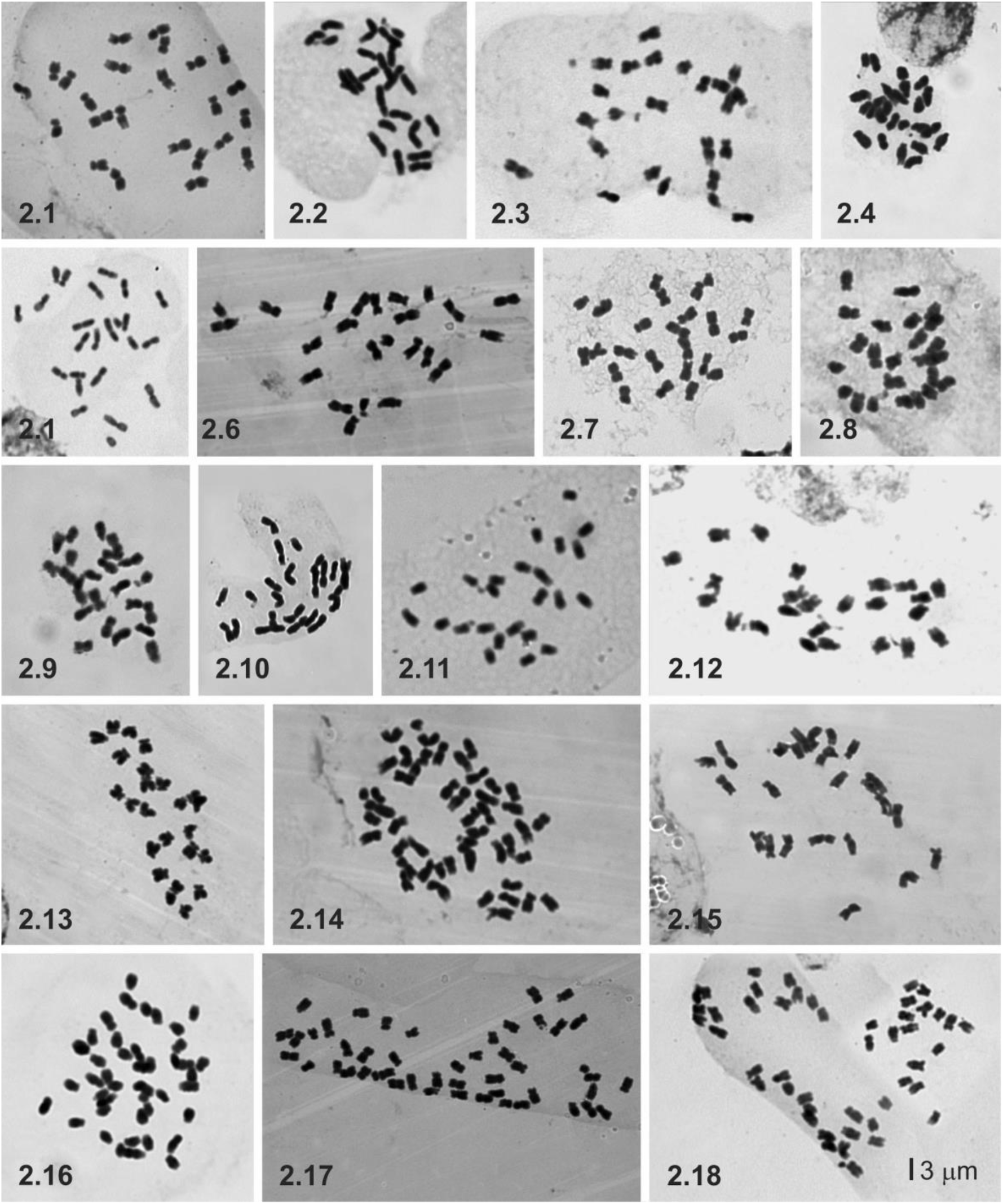
Somatic metaphases of Physalidineae with Giemsa staining. (2.1) *Alkekengi officinarum*. (2.2) *P. hederifolia*. (2.3) *P. ixocarpa*. (2.4) *P. porrecta*. (2.5) *P. cinerascens* var. *spathulifolia*. (2.6) *P. chenopodifolia*. (2.7) *P. cinerascens* var. *cinerascens*. (2.8) *P. philadelphica*. (2.9) *P. fendleri*. (2.10) *P. longifolia*. (2.11) *P. victoriana*. (2.12) *P. virginiana*. (2.13) *P. lagascae*. (2.14) *P. pubescens*. (2.15) *P. viscosa*. (2.16) *Quincula lobata*. (2.17) *P. peruviana*. (2.18) *Chamaesaracha coronopus*. All the pictures at the same scale.

The species studied here were heterogeneous concerning chromosome size, with total average chromosome length (c) between 1.62–4.20 μm (Table 2), although the highest c reported was *Calliphysalis carpenteri* with 4.85 μm (Menzel, 1951; Suppl. mat. S2). The smallest c was found in *P. victoriana* (1.62 μm) and the largest in *P. cinerascens* var. *spathulifolia* (4.20 μm). Accordingly, the largest absolute chromosome length was found in *P. cinerascens* var. *spathulifolia* (5.72 μm) and the smallest in *P. peruviana* (1.08 μm). Among the species analysed, HKL had a 3.11-fold variation (from 19.39 μm in the diploid *P. victoriana* to 60.27 μm in the polyploid *P. pubescens*).

The karyotypes of the Physalidinae studied are remarkably asymmetric, composed of metacentric, submetacentric and subtelocentric chromosomes, with the exception of *P. philadelphica* that only has metacentric and submetacentric chromosomes (Table 1, Fig. 2). Furthermore, notable differences were found among the species studied here in intrachromosomal asymmetry (A_1_ varied from 0.28 to 0.59; Suppl. mat. S3) and also along the chromosome size of the same complement (A_2_ from 0.11 to 0.21; Suppl. mat. S4). On average, arm ratio (r) was 2.12 (range = 1.46-2.77; Table 2; Suppl. mat. S5).

Through the integrative analyses of A_1_ vs A_2_, we resolved that most species are grouped into two large groups. On the one hand, we have the most symmetrical genera such as all the belonging to Iochrominae subtribe (e.g., *Iochroma* Benth.*, Saracha* Ruiz & Pav.), *Witheringia* L'Hér. and *Schraderanthus* Averett. On the other hand, we find *Physalis*, *Chamaesaracha* and *Quincula* grouped for being the most asymmetric genera in the tribe (Fig. 3).The complements of the clade *Physalis* + *Quincula* + *Chamaesaracha* (PQC, from now on) are markedly more asymmetric than those of the other genera of the subtribe Physalidinae (Fig. 3, Suppl. mat. S4-S5), presenting: 1 to 8 m chromosomes (x̄ = 5); 1 to 9 sm chromosomes (x̄ = 4); 1 to 5 st chromosomes (x̄ = 2); t = 1 pair in two species (Table 2, Fig. 2, Suppl. mat. S1).

**Fig. 3.**
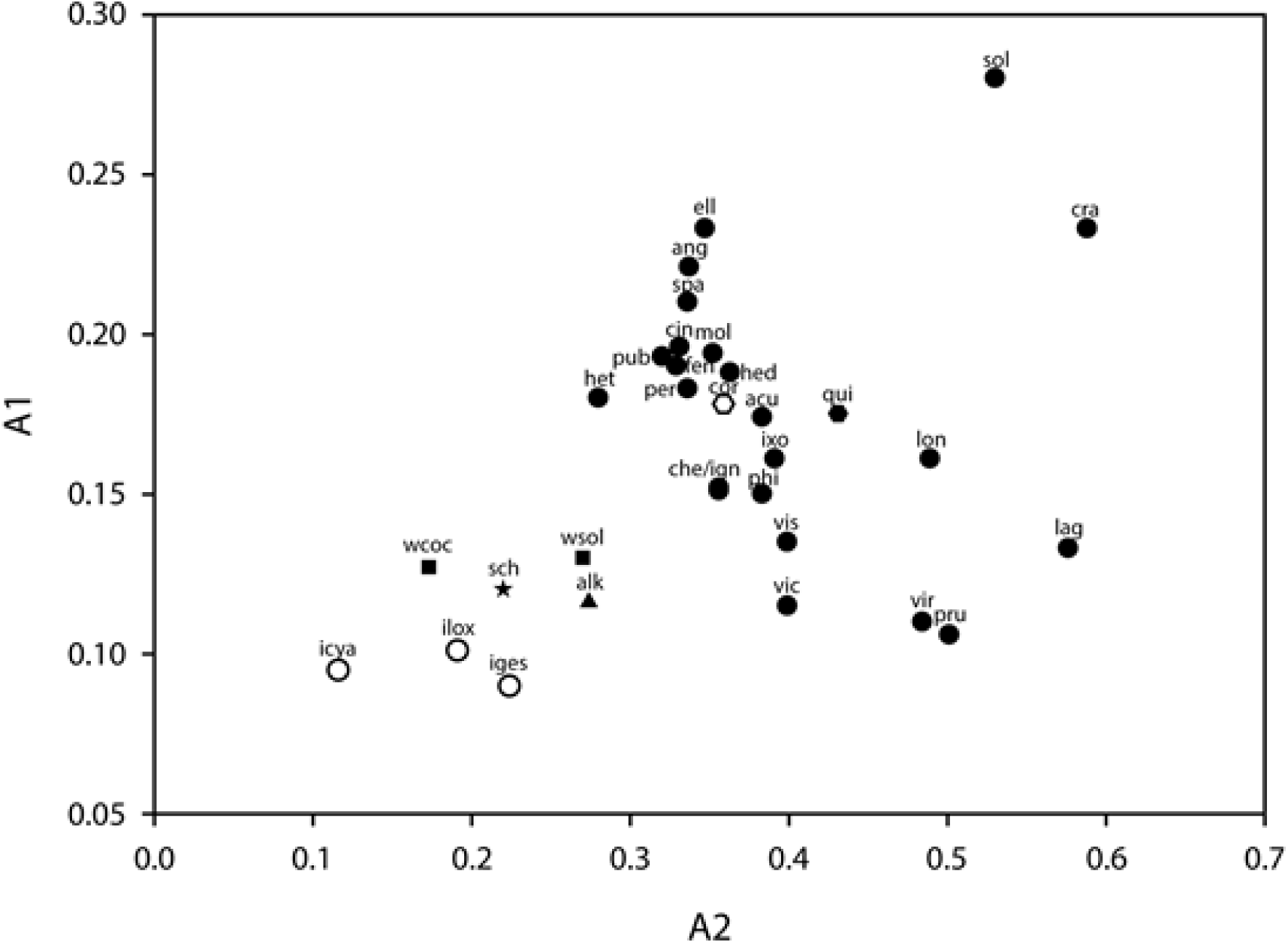
Diagram showing the intrachromosomal asymmetry index (A_1_) plotted against the interchromosomal asymmetry index (A_2_). Species codes are given in table 1. The genera were represented with the following shapes. Solid circles (•) for *Physalis*, empty circles (○) for *Iochroma*, solid triangles (Δ) for *Alkekengi*, solid hexagons (⬡) for *Quincula*, empty hexagons () for *Chamaesaracha*, solid squares (■) for *Witheringia* and a star (*) for *Schraderanthus*.

### 3.2. Ancestral states reconstruction

As a result of the chromosome number reconstructions using ChromEvol, the ‘CONST_RATE’ model (AIC = 65.52) was selected as the best-fitting model for the dataset. Reconstructions supported diploidy (2n = 24) as the ancestral state for the Physalidinae subtribe, and five to six polyploidy events were estimated, plus one aneuploidy event in the monotypic genus *Quincula* (Fig. 4).

**Fig. 4.**
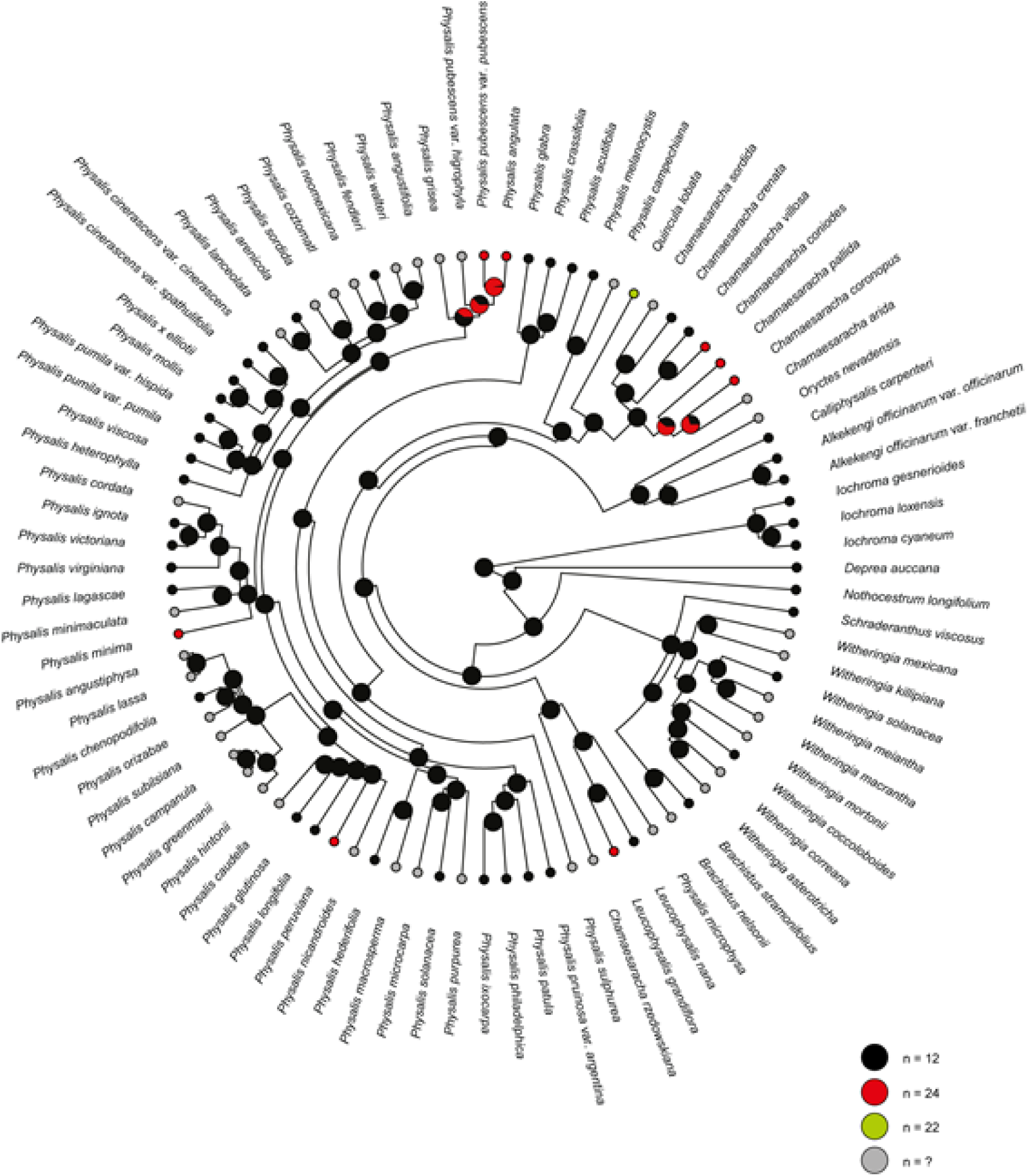
Chromosome haploid number (n) reconstruction in the Physalidinae subtribe with ChromEvol in RASP. Pies at nodes represent frequencies of node states across 10000 simulations of character evolution. Grey pies at tips represent unknown data.

Continuous characters mapped on the MCC tree are represented with phenograms (Fig. 5). The arm ratio (r) clearly differs among clades, with an estimated value around 1.96 for the ancestor of all Physalidinae (Fig. 5.3, Suppl. mat. S5). Accordingly, both asymmetry indices showed strong phylogenetic patterns among different clades (Figs. 5.1 and 5.2, Table 4).

**Fig. 5.**
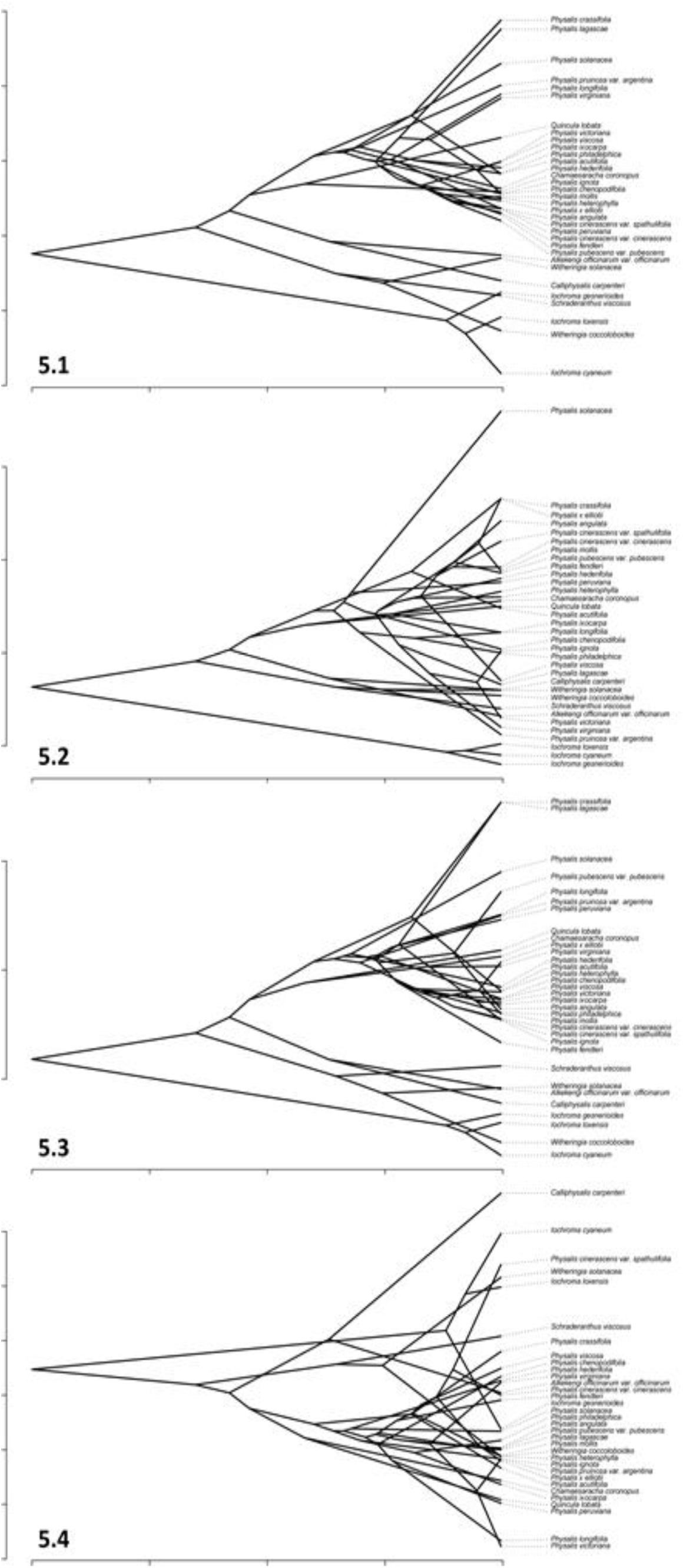
Traitgrams for continuous traits analysed. (5.1) Intrachromosomal asymmetry index (A_1_); (5.2) Interchromosomal asymmetry index (A_2_), (5.3) Mean arm ratio (r); and (5.4) Total average chromosome length (c).

Regarding the Bayesian stochastic mapping for number of submetacentric and subtelocentric chromosomes, both reconstructions resulted in a similar number of changes (23 total shifts in submetacentric chromosome states and 22 total changes for the subtelocentric chromosomes; Table 3). They showed similarities in the number of gains; six gains of four or more pairs of submetacentric chromosomes, and 15 gains of two or more pairs of subtelocentric chromosomes (Table 3; Fig. 6; Suppl. mat. S6).

**Table 3.** Summary of the Stochastic Character Mapping for discrete chromosome features. (TC) mean number of total changes; (C) median number of changes per transition; (95% CI) 95% credibility interval of number of changes. Most frequent transitions are in bold.

| Trait | Model | Character states | TC | TC (95% CI) | Transition | C | C (95% CI) | State at the Physalidinae root |
| --- | --- | --- | --- | --- | --- | --- | --- | --- |
| Submetacentric chromosomes | ARD | 0 = up to three pairs | 23 | 12.40 – 36.39 | 0 > 1 | 4 | 0 – 10.00 | 0 |
|  |  | 1 = between four and six pairs |  |  | 0 > 2 | 1 | 0 – 5.23 |  |
|  |  | 2 = seven or more pairs |  |  | **1 > 0** | 10 | 1.93 – 21.05 |  |
|  |  |  |  |  | 1 > 2 | 6 | 1.80 – 10.72 |  |
|  |  |  |  |  | 2 > 0 | 1 | 0 – 6.82 |  |
|  |  |  |  |  | 2 > 1 | 1 | 0 – 3.71 |  |
| Subtelocentric chromosomes | ARD | 0 = absence | 22 | 12.97 – 31.28 | 0 > 1 | 3 | 0 – 5.70 | 0 |
|  |  | 1 = one pair |  |  | 0 > 2 | 1 | 0 – 3.26 |  |
|  |  | 2 = two or more pairs |  |  | 1 > 0 | 0 | 0 – 7.37 |  |
|  |  |  |  |  | **1 > 2** | 15 | 0 – 21.13 |  |
|  |  |  |  |  | 2 > 0 | 0 | 0 – 6.61 |  |
|  |  |  |  |  | 2 > 1 | 3 | 0 – 16.69 |  |

**Fig. 6.**
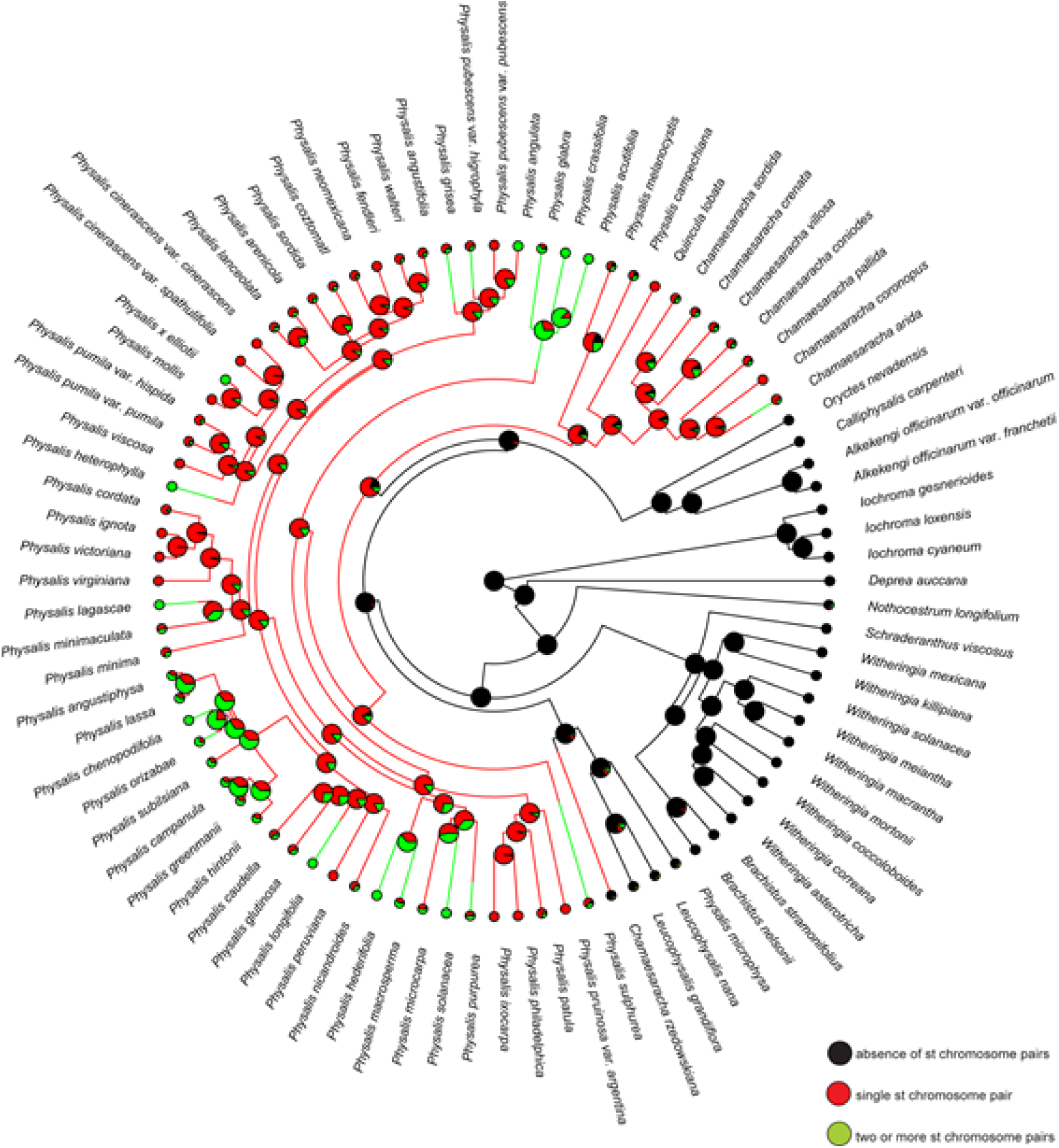
Ancestral state reconstruction of the number of subtelocentric (st) chromosome pairs in the Physalidinae subtribe on the combined MCC tree, using stochastic mapping. Pies at nodes indicate frequencies of node states across 1000 simulations of character evolution and the colours of the tip labels represent tip states.

### 3.3. Phylogenetic signal

Regarding chromosome size (c), the Blomberg’s K was not significantly different from zero (Table 4). For the remaining traits analysed (relationship between arms and intra- and interchromosomal asymmetry), Blomberg’s K was significantly different from zero, but not significantly different from one, indicating a phylogenetic signal in the asymmetry pattern (Table 4). Between both asymmetry indices, the strongest phylogenetic signal was found in the intrachromosomal asymmetry index (A_1_, Table 4).

**Table 4.** Summary of phylogenetic signal (Blomberg’s K) for continuous chromosome traits. PICs: phylogenetically independent contrasts relative to tip shuffling randomization. P-values indicate whether the K-value is significantly different from zero (no phylogenetic signal) and/or from one (signal expected under Brownian Motion). P-values less than 0.05 are bolded.

| Trait | $\lambda$ (Pagel, 1999) | Blomberg's K | P-value of observed vs. random variance of PICs P-value of observed vs. random variance of PICs | P-value of observed vs. random variance of PICs P-value of observed vs. random variance of PICs |
| --- | --- | --- | --- | --- |
|  |  |  | (< 0.05 means K significantly different to zero) | (< 0.05 means K significantly different to one) |
| A1 | 0.99 | 1.23 | **0.0001** | 0.74 |
| A2 | 0.94 | 0.89 | **0.0005** | 0.86 |
| r | 0.91 | 1.02 | **0.0001** | 0.98 |
| c | 0.42 | 0.52 | 0.06 | 0.20 |

## 4. Discussion

### Chromosome number

Solanaceae shows a dysploid series from x=7 to x=14 (cf. Goldblatt & Johnson, 1979 onwards; Rice et al., 2015). A hypothesis on chromosome number changes is still lacking, and even the original basic number for the family is a matter to be clarified. Based on phylogenetic studies, Olmstead et al. (2008) suggested that x = 12 is apomorphic. This is the most frequent base number and characterizes an entire clade, the so-called x = 12 clade, in which Physalideae is placed. Within the x=12 clade, evidence for dysploid changes to x = 13 via translocations has been reported in *Capsicum* L. (Moscone et al., 2006) and *Solanum* L. (subg. *Lycopersicon*; Banks, 1984). However, such type of chromosome change is rare; most species with a chromosome number different from x=12 belong to *Capsicum*, whereas in *Solanum* dysploidy (with x=23) seems to be a synapomorphy of a few species (Chiarini, Sazatornil & Bernardello, 2018). *Quincula lobata* has both diploid and tetraploid races of the basic number 11, with at least occasional aneuploid individuals (2n = 20, Menzel, 1950). Thus, the number x = 11 found in *Quincula* is peculiar, being the Robertsonian translocation the most likely explanation for its origin. Following such a scheme, two telocentric chromosomes would have exchanged their arms, giving as result a big metacentric chromosome and a tiny one formed by the two short arms, which is subsequently lost. An alternative explanation would be the fusion of two telocentric by their short arms with a later inactivation of one centromere. Both explanations are plausible considering that the presence of several pairs of telocentric chromosomes is one of the outstanding features found in the Physalidinae karyotypes.

Except for *Quincula*, the basic number X = 12 is constant in all Physalidinae as far as we know, although the occurrence of several independent events of polyploidy within the group was verified. This can be related to processes of speciation and colonization of new habitats. The role of genome duplications during the invasion of new habitats has been the subject of intense debate (Stebbins, 1985; Soltis et al. 2004). Polyploids frequently have a wider geographical range than their diploid parents (*e.g.*, Schönswetter et al., 2007; Whittemore & Olsen, 2011), probably because they are preadapted to habitats and resources off limits to their parents (Levin, 2003). They have a diversity of alleles that can confer a greater ecological niche than that of diploid progenitors (Brown, 1984; Pound et al., 2004). To this respect, we found two situations in Physalidinae: on the one hand, the genus *Chamaesaracha*, where polyploidy might be related to the colonization of the deserts of Northern Mexico and South Western USA, beyond a hypothetical wetter range of origin. According to Zamora Tavares et al. (2016), the diversification of *Physalis* took place mainly in the Mexican Transition Zone 6.5 mya, when the formation of mountain ranges in this region generated altitudinal and climatic changes that led to the diversification and radiation of taxa. The ancestral distribution for *Chamaesaracha* is the Nearctic and it is characterized by their adaptation to xeric conditions and present more recent divergence dates (5.81–1.59 mya) that coincide with the aridification of North America during the Pliocene (Zamora Tavares et al., 2016). On the other hand, we also found polyploidy in some *Physalis*, which are precisely the species with a wide distribution, probably related to anthropic dispersion. As polyploids often exhibit increased vigour and, sometimes, outmatch their diploid relatives in several aspects, this superiority has been targeted by humans, who have induced polyploidy or have selected natural polyploids in order to obtain improved cultivars (Sattler, Carvalho & Clarindo, 2016). This would be the case of *P. angulata* and *P. pubescens* (growing from USA to Argentina and introduced in the Old World; Yamazaki, 1993; Toledo & Barboza, 2013) and *P. peruviana* (native from Peru but introduced also in the Old World; Brako & Zarucchi, 1993; Zhang et al., 1994).

### Chromosome size

Chromosome length is useful to single out individuals, samples, populations or species, besides being an indirect indicator of the total DNA amount. Solanaceae show notable variation in chromosome size, with an 8-9-fold range (Badr et al., 1997; Chiarini et al., 2018). Most species have small or medium-sized chromosomes with respect to other angiosperms (Levin & Funderburg, 1979; Weiss Schneeweiss & Schneeweiss, 2013), mostly within the 1–3 μm range. Our records on *Physalis* fit in this pattern and confirm previous data (Menzel, 1950, 1951; Rodriguez & Bueno, 2006). We found a 1-fold size variation, similar to the measurements reported by Menzel (1950, 1951), which is still considered a slight variation compared to the sizes found in the family.

Stebbins (1971) proposed a relationship between habit and chromosome size, with perennial species having small chromosomes. Species of subtribe Physalidinae are mostly herbs, whereas its sister clade, Iochrominae, is characterized by the woody habit. However, no marked differences in chromosome size were found when comparing both clades. Also in other woody genera of the family, chromosomes are either small (e.g., *Lycium* L., Stiefkens & Bernardello, 2000; *Lycianthes* (Dunal) Hassl., Acosta, Bernardello, Guerra & Moscone, 2005), or medium-sized (*Latua pubiflora* (Griseb.) Baill., Chiarini et al., 2010).

Genome downsizing is a phenomenon that apparently occurs in polyploids (Leitch & Bennett, 2004). Theoretically, polyploids are expected to have larger DNA content than their diploid progenitors, increasing in direct proportion with ploidy, but this is true mostly in newly formed polyploid series. Contrarily, there are cases in which, either the mean 1C DNA amount does not increase proportionally with ploidy, or the mean DNA amount per basic genome (2C value divided by ploidy) tend to decrease with increasing ploidy (Leitch & Bennett, 2004). Gene reordering and deletion of redundant material have been proposed to account for this DNA content decrease, and these changes may be reflected on chromosome length, considering that length is an indirect indicator of the total DNA amount. Also, polyploids may have proportionally less DNA amount but yet share the same chromosome morphology with their diploid relatives. For instance, Poggio, Realini, Fourastié, García & González (2014) found that in *Hippeastrum* (Amarillydaceae), polyploid species show less DNA content per basic genome than diploid species, but the typical bimodal karyotype is preserved, even in the presence of genome downsizing. The authors suggest that constancy of the karyotype is maintained because changes in DNA amount are proportional to the length of the whole chromosome complement and vary independently in the long and short sets of chromosomes. In the accessions here studied, we found also a constancy: despite they can vary on c, species of the PQC clade have similar formulae, which in average would be 5m + 4sm + 2st + 1t. Even polyploids have formulae which are the double of this average formula, however their chromosomes are slightly smaller compared to the diploid species.

### Karyotype asymmetry

Within the x = 12 clade there are some genera that present a high constancy in chromosome morphology, with symmetrical karyotypes and a majority of m chromosomes of rather similar size, e.g., *Lycium* (Stiefkens & Bernardello, 2000), but at the same time, complements that include st and t chromosomes are found in species of *Nicotiana*, *Capsicum*, *Jaborosa*, in some *Solanum* (Acosta et al., 2005; Chiarini & Barboza, 2008; Menzel, 1951) and in *Physalis* (Menzel, 1951; Venkateswarlu & Raja Rao, 1977, 1979a, 1979b; Rodríguez & Bueno, 2006). Karyotypes that are highly symmetrical have been considered primitive (Stebbins, 1971), but concurrently, a karyotype orthoselection has been hypothesized for the conservation of complements with homogeneous chromosomes (Brandham & Doherty, 1998; Moscone et al., 2003). It is difficult to establish the direction of karyotype evolution, as many reversals might have occurred (Stace, 2000), and karyotype asymmetry might be a transient state rather than a derived evolutionary end (Mandakova & Lysak, 2008). However, our reconstructions of chromosomal characters indicate that the most probable character state for the ancestor of all Physalideae and Physalidinae had a formula with none st chromosomes. The probability of having a karyotype with at least one st becomes higher with the common ancestor of the clade PQC, and this trend increases in some species groups within *Physalis* which present formulae with more than two st chromosomes. Thus, the karyotype changes in Physalideae would be directional towards an asymmetric karyotype in the PQC clade.

Transitions between karyotype formulae can be interpreted as evidence of chromosome rearrangements. Specifically, formulae with st or t chromosomes are seemingly the result of a deletion or translocation of the entire or part of one arm (Weiss Schneeweiss & Schneeweiss, 2013). Species of the PQC clade share markedly asymmetrical formulae (5m +4sm +2st +1st or 6m +4sm +2st). *Witheringia* is symmetrical (9m + 3sm, Barboza et al., 2010; Chiarini et al., 2010) and the rest of genera in the tribe Physalideae are so (*Athenaea* Sendtn. And *Deprea* Raf., 9m + 3sm; Deanna et al., 2014; Chiarini et al., 2017), whereas subtribe Iochrominae is even more symmetrical (11m + 1 sm or 12m, Deanna et al., 2018). The most parsimonious explanation for this situation is that large chromosome rearrangements would have occurred when the common ancestor for PQC separated from the ancestor that gave rise to the clade formed by *Calliphysalis* + *Alkekengi* + *Oryctes*, this latter retaining a symmetrical karyotype. This is another argument that supports the segregation of these three genera outside of *Physalis*, where they were previously assigned (Pretz & Deanna, 2020).

According to our ASR, symmetrical karyotypes are plesiomorphic, being asymmetry a synapomorphy of the PQC clade, while *Calliphysalis* + *Alkekengi* + *Oryctes* would have conserved symmetry. This supports the idea that shifts in chromosome morphology are quite common when considering wide frames of time. The papers of Wu & Tanskley (2010) and Chiarini et al. (2018) provided estimates of such frequency by dividing evolutionary time by number of chromosome rearrangements (what yields one rearrangement per complement approximately every 2.93 my, or 0.34 rearrangement per million years). This supports the idea that diversification in the PQC clade has been recent, and not enough time has passed for so many rearrangements to accumulate and blur this uniform pattern of asymmetric karyotypes. Even though, chromosome traits are informative to distinguish clades: from the traitgrams of Fig. 5 is clear that the A_1_ index has a strong phylogenetic signal, separating Iochrominae from the PQC clade. This congruence between karyotypes evolution and the splits and differentiation of clades within phylogenies has already noticed in genera from different plant families (*e.g.*, Blöch et al., 2009; Baltisberger & Hörandl, 2016). It is unlikely that extant species with divergent karyotypes are able to cross, whereas species with the same karyotype should be capable of producing hybrids.

### Taxonomic implications

Our results provide evidence on the following taxonomic issues: 1) Support to differentiate the genus *Quincula* from *Physalis* and *Chamaesaracha*, on the grounds of its exclusive basic number and particular morphology (Barboza, 2000); 2) Maintenance of *P. solanacea* within *Physalis* instead of placing it in a monotypic genus (as *Margaranthus solanaceus* Schltdl.), based on similarities in formulae and asymmetry indices as well as phylogenetic placement (Whitson & Manos, 2005; Zamora-Tavares et al., 2016; Deanna et al., 2019); 3) Segregation of *Calliphysalis* Whitson and *Alkekengi* Mill. out of *Physalis*, on the basis of their symmetrical karyotypes, morphological features and phylogenetic placement (*cfr.* Pretz & Deanna, 2020); and 4) Support to distinguish *Deprea* from *Physalis*: some species of *Deprea* have inflated fruiting calyx which has confused them to be mistaken by *Physalis*, but both genera can be easily distinguished by their karyotypes, as well as phylogenetic distance and floral traits (Whitson & Manos, 2005; Deanna et al., 2019).

However, to complete the chromosome evolution picture within Physalideae and confirm that asymmetry is a characteristic of only the PQC clade, more karyotype studies are needed in the remaining clades of the tribe (especially those fromAsia, like *Physaliastrum* Makino; Zhang et al., 1994). Chromosome number and morphology are still unknown for the genera *Mellissia* Hook.f., *Cuatresia* Hunz., *Brachistus* Miers and also for the genera recently segregated from *Physalis*: *Tzeltalia* E.Estrada & M.Martínez, *Archiphysalis* Kuang, and *Darcyanthus* Hunz. ex N.A.Harriman (Pretz & Deanna, 2020).

### Conclusions

In accordance with our results on ASR, the common ancestor to all the subtribe Physalidinae was a diploid with 2n=24, with a karyotype with up to three sm chromosomes, an arm ratio of ca. 1.96, and approximately A_1_ = 0.35 and A_2_ = 0.14. Polyploidy occurred six times after the divergence of Physalidinae from the rest of the Physalideae, and one aneuploidy event was involved in the split of *Quincula*. These numeric alterations can be related to processes of speciation and colonization of new habitats. The main evolutionary trends with respect to the ancestor were: an increase of asymmetry in the lineage that originated the PQC clade; a decrease of asymmetry in Iochrominae and *Witheringia*. Both asymmetry indices allow to differentiate genera within the subtribe. Overall, our study suggests that Physalideae diversification has been accompanied by karyotype changes, which can be applied in the delimitation of genera within the group.

The results here analysed raise questions for future prospects: 1) Is there a relationship among karyotype formulae and different macromorphological and ecological features, as habit, niche or life form? Such relationships are usually hypothesized but not tested. For example, Menzel (1950) proposed a correlation among ploidy, seed size, and geographic range, suggesting that individuals with larger seeds would have higher ploidy, but this matter remains inconclusive. 2) Concerning traits of agronomic value, do the polyploid Physalidinae outmatch in size, vigour or even environmental resilience their diploid parents? The most important crops worldwide are polyploids (Sattler et al., 2016) so it would be useful to address this question for the ongoing improvement programs on ground cherries and golden berries (*Physalis*).

## Supporting information

Supplemental mat S1-S6

## 5. Acknowledgements

The authors thank the National Council for Scientific Research and Techniques (CONICET), National Agency for Scientific Promotion and Technological (FONCyT, grant # PICT 2017-2370 and 2016-1525) and SECyT (National University of Córdoba, Argentina, grant # 203/14) for financial support, equipment and facilities. We thank Chelsea Pretz, Tiina Sarkinen, Sandra Knapp from http://solanaceaesource.org/, and G. Chaniot from https://www.inaturalist.org/ for their contribution in photographic material.

