## Supplemental mat S1-S6 for "Karyotype asymmetry shapes diversity within the physaloids (Physalidinae, Physalideae, Solanaceae)"

**APPENDIX S1.** Idiograms of *Alkekengi*, *Chamaesaracha*, *Physalis* and *Quincula* species.

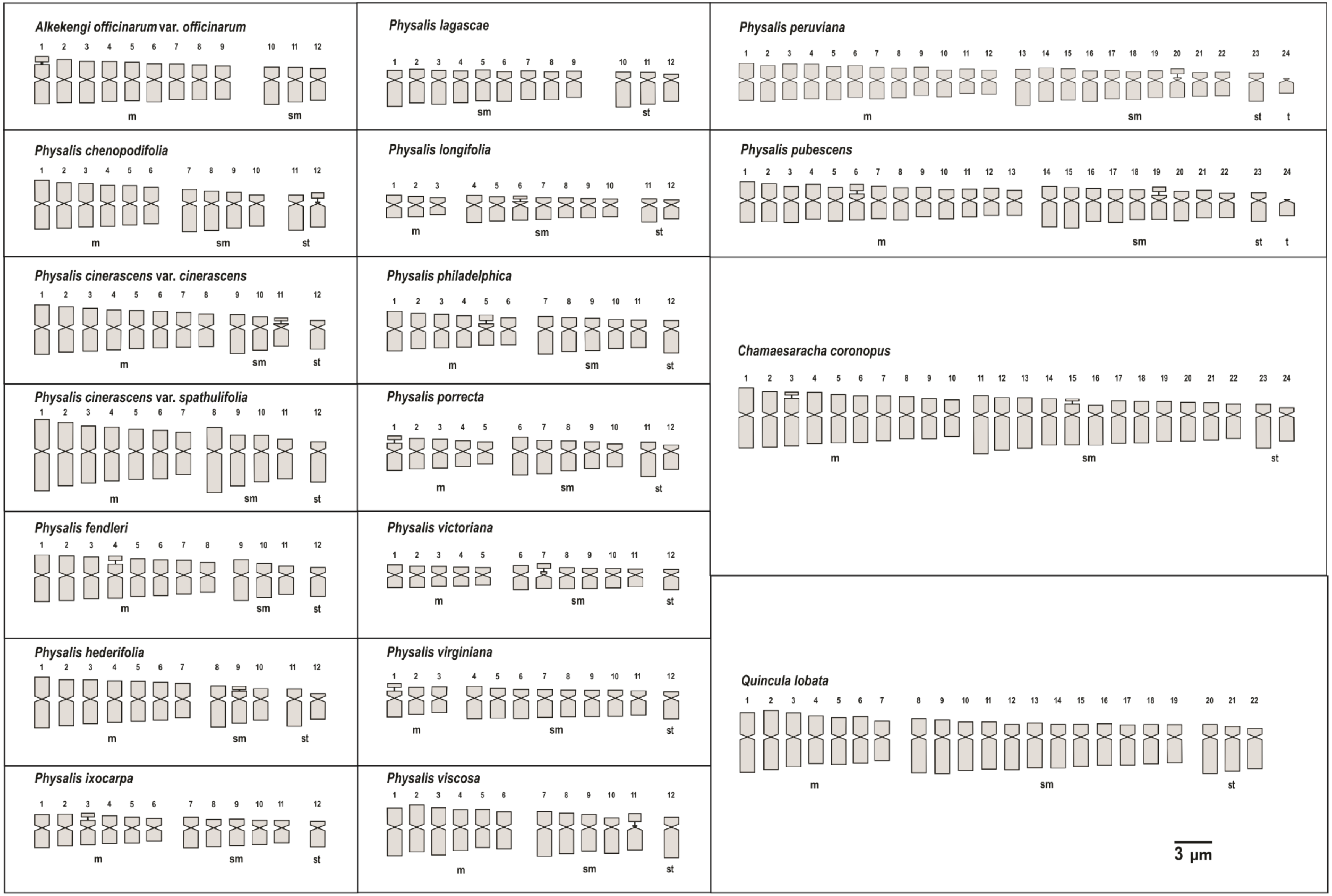

**APPENDIX S2.** Maximum likelihood reconstruction of total average chromosome length (c) values on the combined MCC tree.

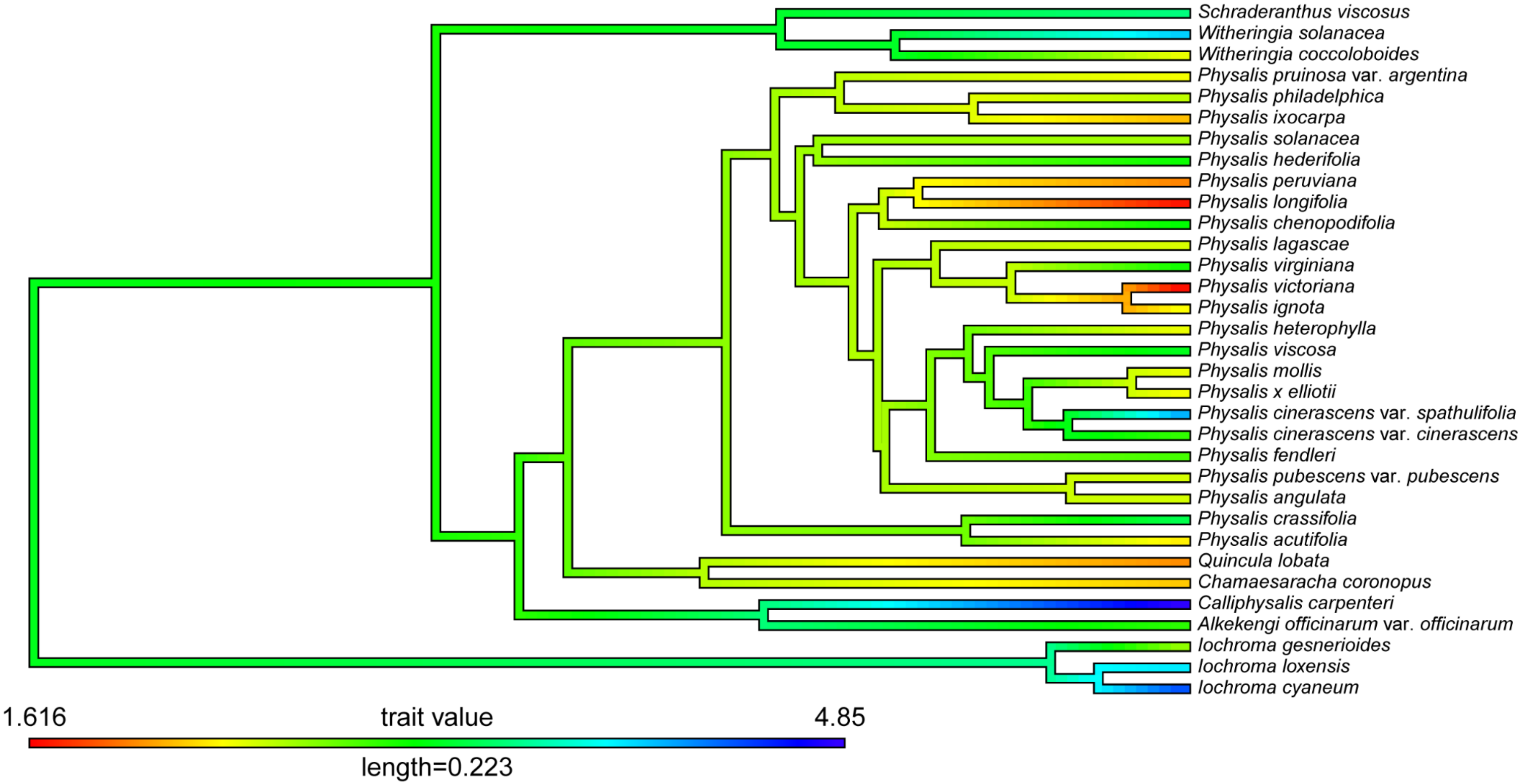

**APPENDIX S3.** Maximum likelihood reconstruction of intrachromosomal asymmetry index ($A_1$) values on the combined MCC tree.

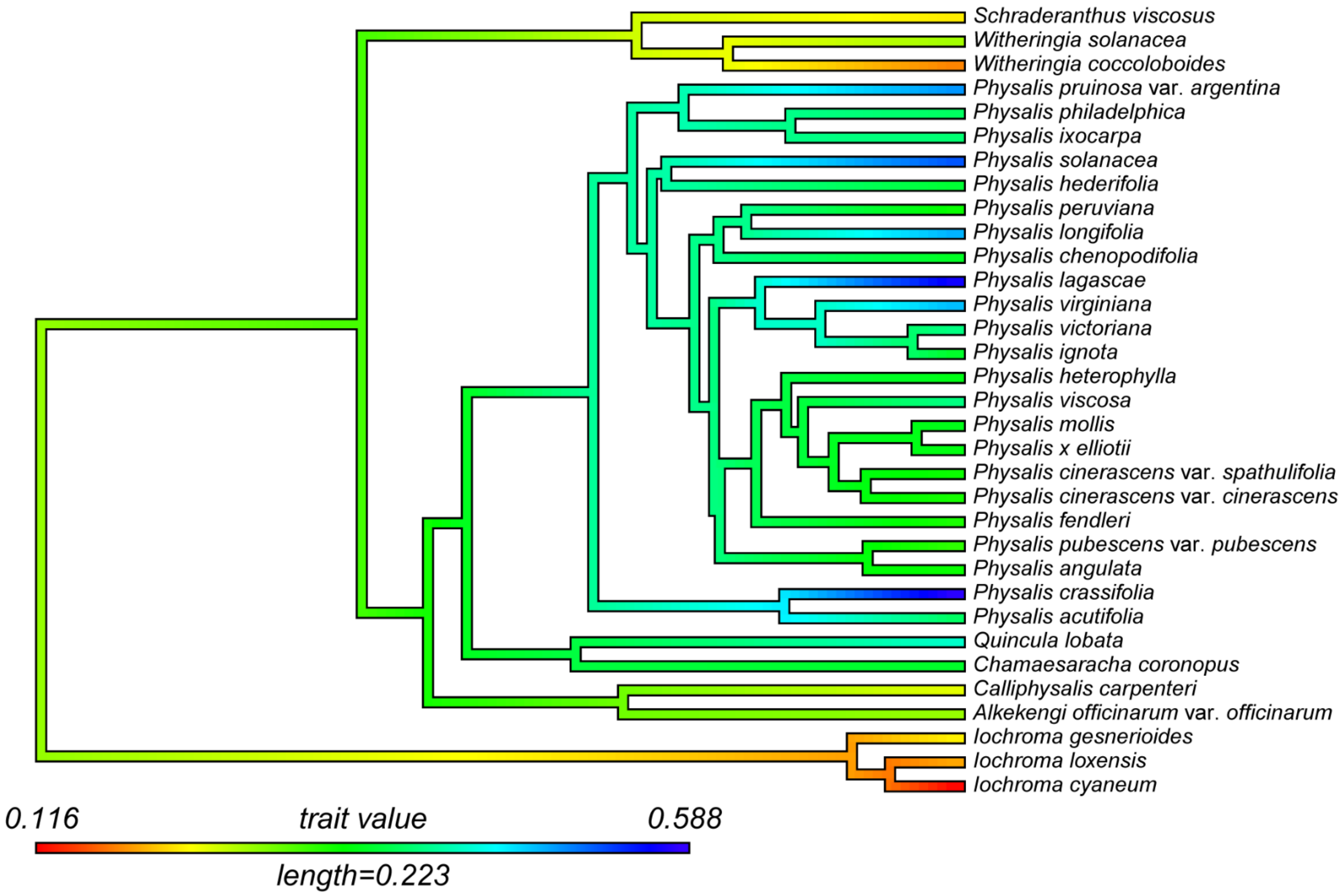

**APPENDIX S4.** Maximum likelihood reconstruction of interchromosomal asymmetry index ($A_2$) values on the combined MCC tree.

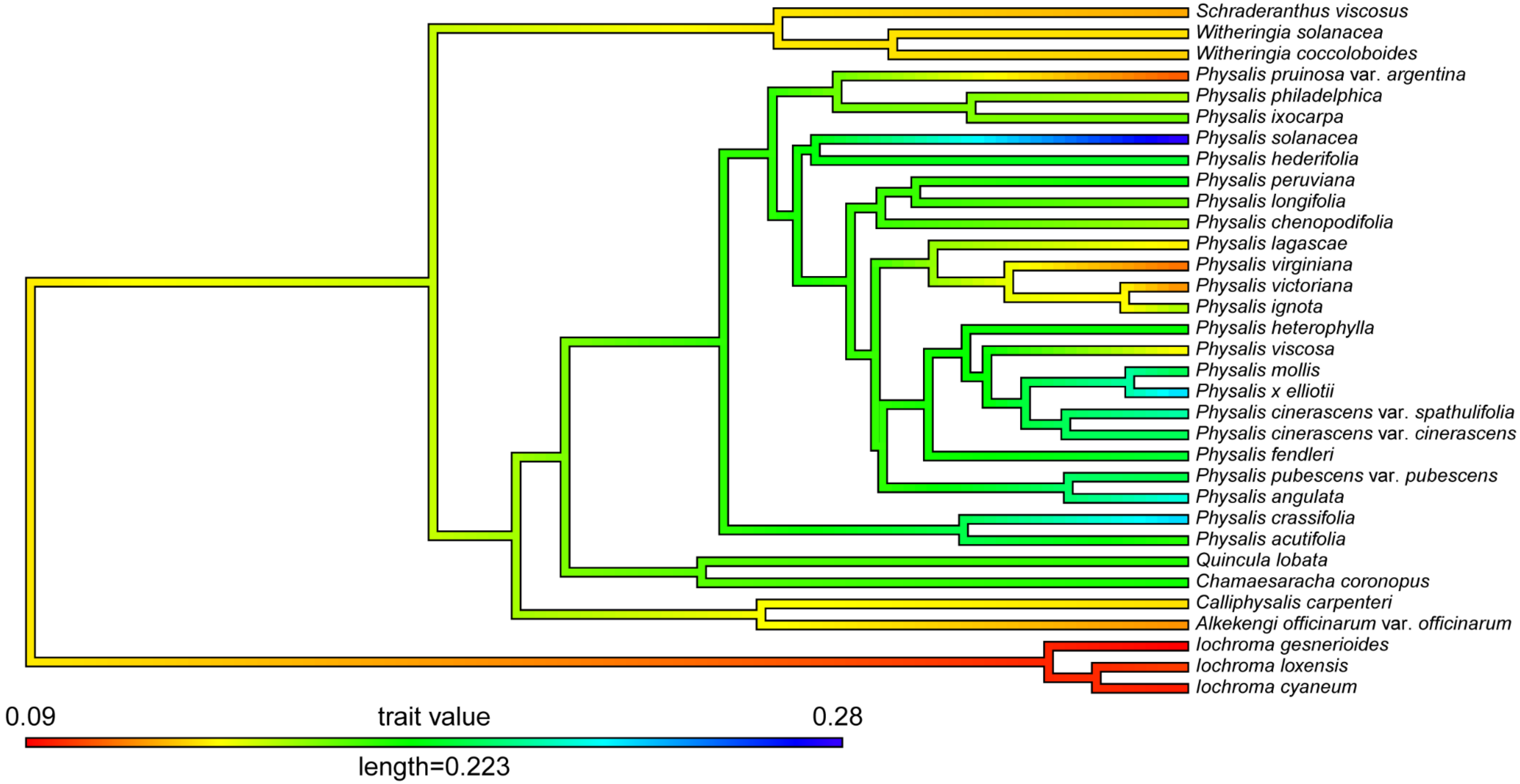

**APPENDIX S5.** Maximum likelihood reconstruction of mean arm ratio (r) values on the combined MCC tree.

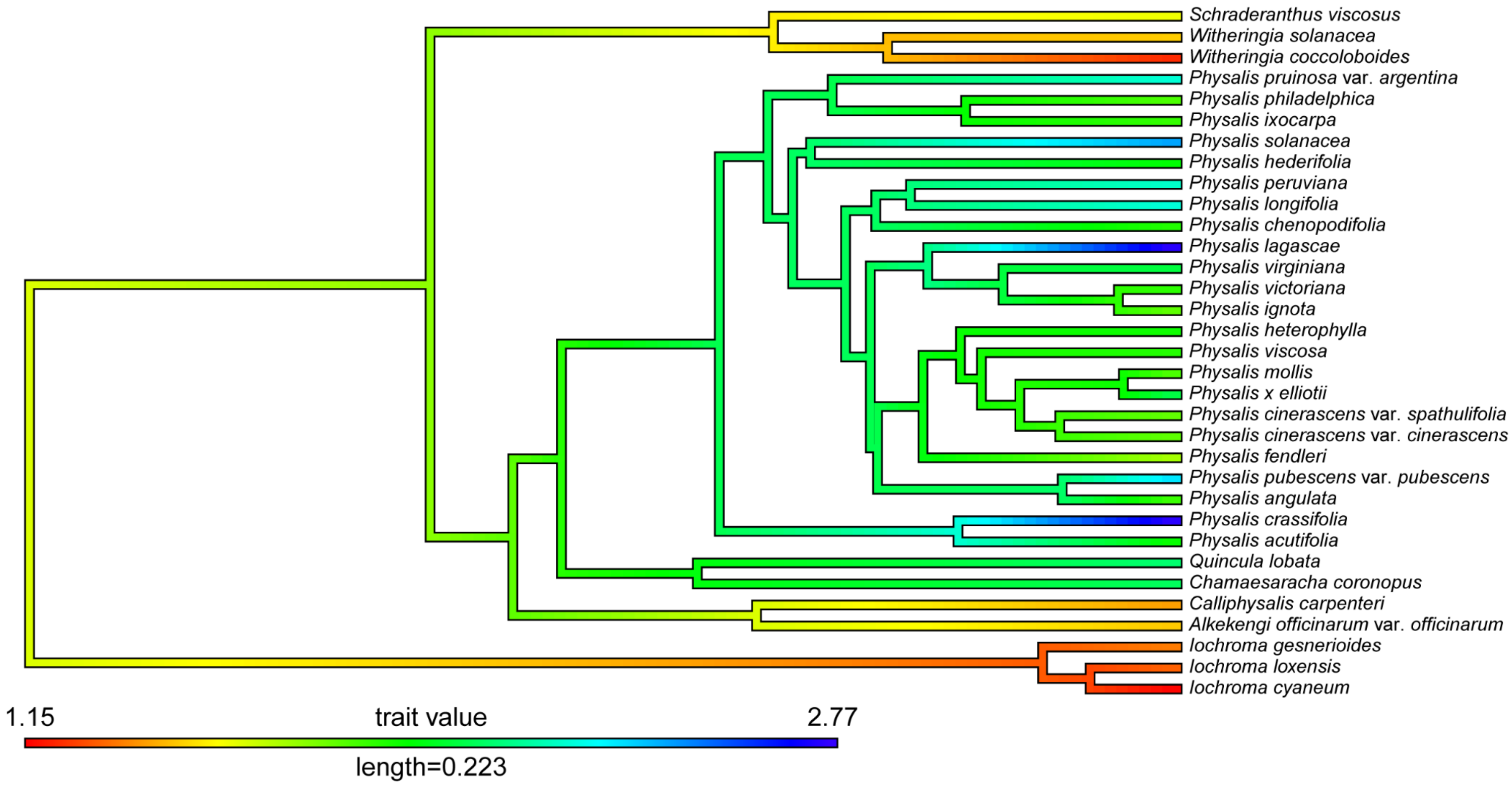

**APPENDIX S6.** Ancestral state reconstruction of the number of submetacentric (sm) chromosome pairs in the Physalidinae subtribe on the combined MCC tree, using stochastic mapping. Pies at nodes indicate frequencies of node states across 1000 simulations of character evolution and the colours of the tip labels represent tip states.

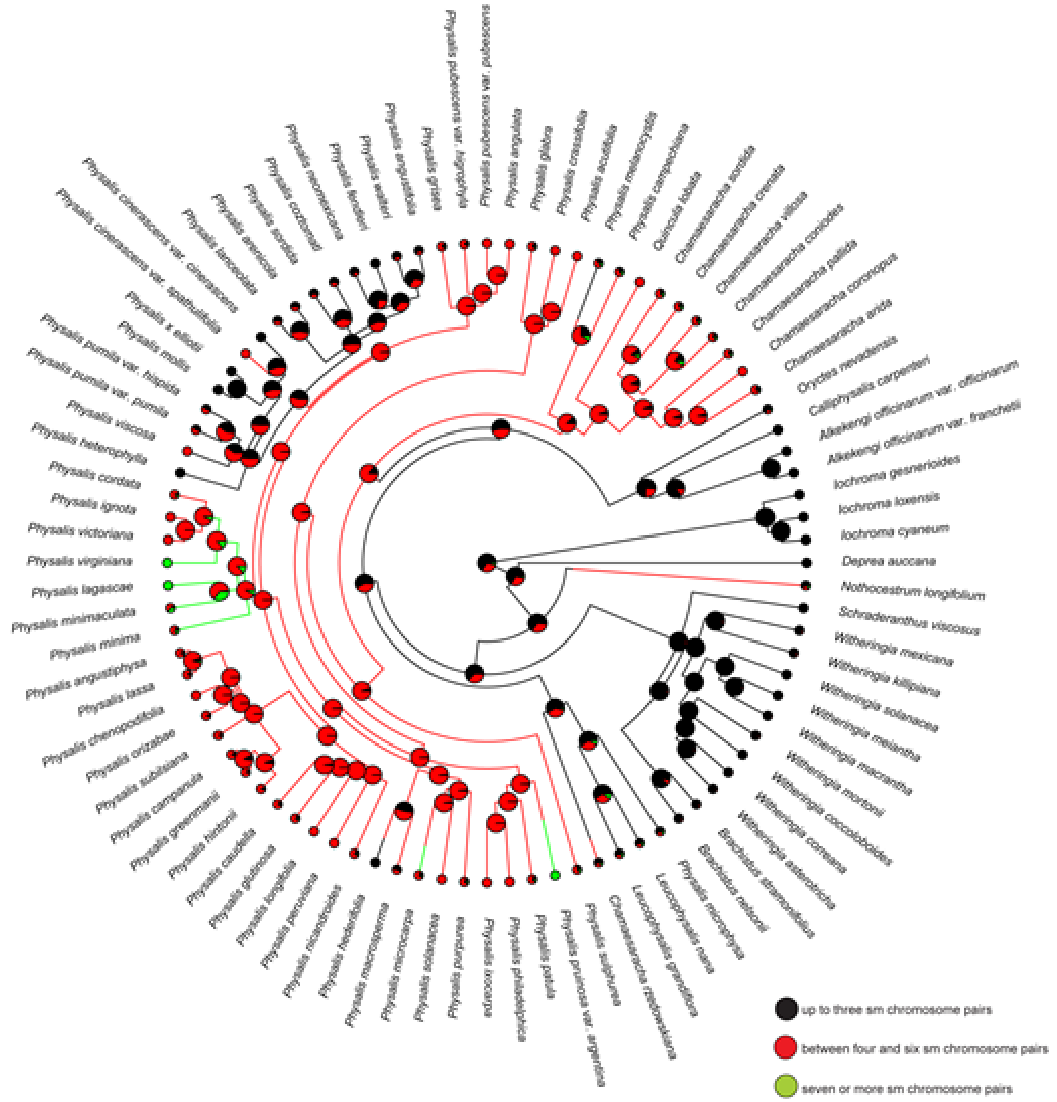
